# Whole-brain dynamics in aging: disruptions in functional connectivity and the role of the rich club

**DOI:** 10.1101/2020.06.29.164343

**Authors:** Anira Escrichs, Carles Biarnes, Josep Garre-Olmo, José Manuel Fernández-Real, Rafel Ramos, Reinald Pamplona, Ramon Brugada, Joaquin Serena, Lluís Ramió-Torrentà, Gabriel Coll-De-Tuero, Luís Gallart, Jordi Barretina, Joan C. Vilanova, Jordi Mayneris-Perxachs, Marco Essig, Chase R. Figley, Salvador Pedraza, Josep Puig, Gustavo Deco

## Abstract

Normal aging causes disruptions in the brain that can lead to cognitive decline. Resting-state fMRI studies have found significant age-related alterations in functional connectivity across various networks. Nevertheless, most of the studies have focused mainly on static functional connectivity. Studying the dynamics of resting-state brain activity across the whole-brain functional network can provide a better characterization of age-related changes. Here we employed two data-driven whole-brain approaches based on the phase synchronization of blood-oxygen-level-dependent (BOLD) signals to analyze resting-state fMRI data from 620 subjects divided into two groups (‘middle-age group’ (n=310); age range, 50-65 years vs. ‘older group’ (n=310); age range, 66-91 years). Applying the Intrinsic-Ignition Framework to assess the effect of spontaneous local activation events on local-global integration, we found that the older group showed higher intrinsic ignition across the whole-brain functional network, but lower metastability. Using Leading Eigenvector Dynamics Analysis, we found that the older group showed reduced ability to access a metastable substate that closely overlaps with the so-called rich club. These findings suggest that functional whole-brain dynamics are altered in aging, probably due to a deficiency in a metastable substate that is key for efficient global communication in the brain.

## Introduction

Normal aging is associated with changes in the structure and function of the brain that could lead to cognitive decline and worse quality of life (Li et al., 2015). Studying the mechanisms of brain aging may identify interventions to prevent or slow age-related deterioration and improve our understanding of the mechanisms involved in neurodegenerative diseases (Ferreira and Busatto, 2013). In recent years, noninvasive resting-state functional magnetic resonance imaging (fMRI) paradigms from spontaneous blood-oxygen-level-dependent (BOLD) signals have proven useful in studying age-related changes in brain function (Ystad et al., 2011). Resting-state fMRI shows coherent spontaneous low-frequency fluctuations across brain regions and the organization of these regions into different functional networks (Zuo et al., 2010). Studies of functional connectivity have suggested age-related alterations in different resting-state networks (Raichle et al., 2001; Wang et al., 2010; Ferreira and Busatto, 2013; Grady et al., 2016), even in cognitively preserved older adults (Damoiseaux et al., 2008; Onoda et al., 2012). Other studies (Li et al., 2015; Grady et al., 2016; Fjell et al., 2017) have suggested that overactivation in functional connectivity across resting-state networks may be related to compensatory mechanisms.

Although functional connectivity studies have demonstrated reliable age-related changes, it remains unclear how brain networks cooperate to handle aging-associated declines, especially considering the effects of averaging on measurements of functional connectivity during rest (Hutchison et al., 2013). In this line, growing evidence indicates that functional connectivity among brain networks is not static over time; rather, different brain regions connect and disconnect from one another in highly complex temporal dynamics (Deco et al., 2011; Hutchison et al., 2013; Sporns, 2013; Zalesky et al., 2014; Ponce-Alvarez et al., 2015). In other words, even in the resting state, brain networks fluctuate in response to different contexts or external stimuli. Capturing statistical properties of fMRI data beyond classical static functional connectivity can facilitate the interpretation of brain functioning during the resting scan from new perspectives. This approach assumes that mental operations arise from neural communication involving coherent and flexible oscillatory activity between functional groups of neurons (Hutchison et al., 2013; Deco and Kringelbach, 2016). The term metastability (Deco and Kringelbach, 2016) refers to the temporal variability of the functional connectivity that arises from the underlying structural connectivity (the human connectome) (Sporns et al., 2005). Optimal brain function is thought to occur within a range of metastable patterns that reflects a balance between the synchronization and adaptive reconfiguration of the functional connections among the different regions that make up the structural network (Cabral et al., 2011).

Dynamic (time-varying) functional connectivity has been explored across the lifespan (Nomi et al., 2017), across different states of consciousness (Deco et al., 2017b; Escrichs et al., 2019; Lord et al., 2019), in patients with brain disorders (Puig et al., 2018), and during healthy aging (Tian et al., 2018; Nobukawa et al., 2019). One study that evaluated resting-state fMRI data from 250 subjects to examine patterns of resting-state functional connectivity over time found that dynamic connectivity patterns are consistent across groups (Abrol et al., 2016). Another study (Yin et al., 2016) found that age-related changes in the functional flexibility of the brain differ in different regions of the cerebral cortex. A recent study in 188 cognitively healthy elderly individuals (Lou et al., 2019) found that frequency-specific brain network diversity decreased with increasing age at both the whole-brain and regional levels. Thus, exploring dynamic functional connectivity promises to enrich our knowledge of the functional organization of the brain, but little is known about changes in dynamic functional connectivity during aging.

In this work, we explored age-related changes in dynamic functional connectivity across the whole-brain network, applying two recently developed data-driven methods based on the phase synchronization of resting-state fMRI BOLD signals to a large dataset from healthy human adults. We studied two aspects of whole-brain functional connectivity in middle-aged subjects versus older subjects: (1) the effects of spontaneously occurring local activation events on local-global integration through the intrinsic-ignition framework (Deco and Kringelbach, 2017; Deco et al., 2017b) and (2) recurrent dynamic functional connectivity patterns across time (here, referred to as metastable substates), their duration, and their probability of occurrence through Leading Eigenvector Dynamics Analysis (LEiDA) (Cabral et al., 2017).

## Materials and Methods

### Subjects

The study population was drawn from the 1030 subjects aged ≥50 years who participated in the population-based Aging Imageomics Study (Puig et al., 2020) from whom data were collected between November 2018 and June 2019. We excluded subjects for whom the full brain imaging dataset was unavailable: those who did not undergo the complete brain imaging protocol including fMRI (n=23), those with MRI acquisition errors (n=192), and those with uncorrectable motion artifacts (n=92; see the Preprocessing section below). Thus, the inclusion criteria were met by 723 subjects [310 aged < 65 years (middle-aged group) and 413 aged ≥ 65 years (older group)]. To homogenize the size of the samples in the two groups, we randomly selected 310 subjects from those aged ≥65 years. The middle-aged group comprised 310 subjects aged < 65 years (mean age, 60.2±3.7 y), and the older group comprised 310 subjects aged ≥ 65 years (mean age, 71.8±4.5 y). Table 1 reports details about subjects’ social and physical status. The ethics committee at the Dr. Josep Trueta University Hospital supervising the study approved the study protocol, and all subjects provided written informed consent.

**Table 1.** Demographic and clinical characteristics

|  | **Overall sample**<br>(n=620) | **Middle-age group**<br>**(50-64 years)**<br>(n=310) | **Older group**<br>**(≥ 65years)**<br>(n=310) |
| --- | --- | --- | --- |
| **Sex** (female), n (%) | 307 (49.5) | 169 (54.5) | 138 (44.5) |
| **Age**, mean (SD) | 65.9 (7.2) | 60.2 (3.7) | 71.8 (4.5) |
| **Age groups**, n (%) |  |  |  |
| 50-64 | 310 |  |  |
| 65-91 | 310 |  |  |
| **Education level** *, n (%) |  |  |  |
| No schooling | 18 (2.9) | 2 (0.7) | 16 (5.2) |
| Primary (ISCED 1) | 324 (52.8) | 133 (43.3) | 191 (62.2) |
| Secondary (ISCED 2) | 90 (14.7) | 55 (17.9) | 35 (11.4) |
| Professional (ISCED 3-4) | 107 (17.4) | 67 (21.8) | 40 (13.0) |
| University (ISCED 5-8) | 75 (12.2) | 50 (16.3) | 25 (8.1) |
| **Body mass index** **, n (%) |  |  |  |
| <18.5 kg/m<sup>2</sup> | 5 (0.8) | 5 (1.6) | 0 (0.0) |
| 18.5 kg/m<sup>2</sup>-24.9 kg/m<sup>2</sup> | 156 (25.2) | 96 (31.2) | 60 (19.4) |
| 25.0 kg/m<sup>2</sup>-29.9 kg/m<sup>2</sup> | 279 (45.1) | 118 (38.3) | 161 (51.9) |
| ≥ 30 kg/m<sup>2</sup> | 178 (28.8) | 89 (28.9) | 89 (28.7) |
| **Physical activity groups (IPAQ)**, n (%) † |  |  |  |
| High | 303 (51.5) | 136 (47.2) | 167 (55.7) |
| Moderate | 248 (42.2) | 130 (45.1) | 118 (39.3) |
| Low | 37 (6.3) | 22 (7.6) | 15 (5.0) |
| **Weight** (kg), mean (SD) | 75.6 (14.1) | 75.1 (15.5) | 76.1 (12.5) |
| **Height** (cm), mean (SD) | 164 (9.0) | 164 (9.1) | 163 (9.1) |
| **Systolic arterial pressure** (mmHg), mean (SD) | 138.8 (19.4) | 135.1 (19.2) | 142.6 (18.8) |
| **Diastolic arterial pressure** (mmHg), mean (SD) | 84.1 (10.6) | 85.0 (10.1) | 83.1 (11.0) |
| **Hypertension**, n (%) # | 289 (46.9) | 120 (38.8) | 169 (55.0) |
| **Diabetes mellitus**, n (%) # | 139 (22.5) | 47 (15.2) | 92 (29.9) |
| **Dyslipidemia**, n (%) # | 181 (29.4) | 79 (25.5) | 102 (33.3) |

\* 6 missing values; \*\* 2 missing values; † 32 missing values; # 4 missing values
IPAQ: International Physical Activity Questionnaire

### Image acquisition

Images were acquired on a mobile 1.5T scanner (Vantage Elan, Toshiba Medical Systems at the beginning of the study; now Canon Medical Systems) with an 8-channel phased-array head coil with foam padding to restrict head motion and noise-cancelling headphones. Brain MRI studies included the acquisition of a high-resolution axial T1-weighted sequence (number of slices=112; repetition time (TR)=8 ms; echo time (TE)=4.5 ms; flip angle=15°; field of view (FOV)=235×235 mm; and voxel size=1.3×1.3×2.5 mm) for structural imaging and a gradient echo-planar imaging (EPI) sequence (TR=2500 ms; TE=40 ms; flip angle=83°; FOV=230×230 mm; and voxel size = 3.5×3.5×5 mm without gap) with 122 continuous functional volumes acquired axially for 5 minutes for resting-state fMRI. Subjects were asked to keep their eyes closed, relax, remain as motionless as possible, and not fall asleep.

### Image Preprocessing

T1-weighted and EPI images were automatically oriented using Conn (Whitfield-Gabrieli and Nieto-Castanon, 2012). For preprocessing, we used the Data Processing Assistant for Resting-State fMRI (DPARSF) toolbox [(Chao-Gan and Yu-Feng, 2010), www.rfmri.org/DPARSF], based on Statistical Parametric Mapping (SPM12) (http://www.fil.ion.ucl.ac.uk/spm). Preprocessing included: (1) discarding the first 5 volumes from each scan to allow for signal stabilization; (2) slice-timing correction; (3) realignment for head motion correction across different volumes; (4) co-registration of the functional image to the T1-weighted image; (5) normalization by using T1 image unified segmentation; (6) nuisance covariates regression: six parameters from the head motion correction, the white matter signal, and the cerebrospinal fluid signal using CompCor (Behzadi et al., 2007); (7) removal of the linear trend in the time series; (8) spatial normalization to the Montreal Neurological Institute standard space; (9) spatial smoothing with 6 mm full width at half-maximum Gaussian kernel; and (10) band-pass temporal filtering (0.01-0.025 Hz). We used a cutoff of 0.25 Hz for the maximum detectable frequency in typical resting-state fMRI acquisitions (Yuen et al., 2019). Then, the time series were extracted using a resting-state atlas of 214 brain areas (without the cerebellum) which ensures the functional homogeneity within each brain subunit (Shen et al., 2013; Finn et al., 2015).

We excluded a total of 92 subjects for head rotation or movement (67 for head rotation > 2 mm or 2° and 25 for frame-wise displacement (Jenkinson et al., 2002; Yan et al., 2013), defined as head motion > 2 standard deviations above the group average in > 25% timepoints).

### Phase Synchronization

We computed the instantaneous phase of the BOLD signals between each pair of brain areas at each timepoint. First, to avoid artifacts, we band-pass filtered the BOLD time series within the narrowband (0.04-0.07 Hz) (Glerean et al., 2012) (Figure 1.1A). Then, we obtained the analytic signal, *a*(*t*), of the filtered time series of each brain area by computing the Hilbert transform (HT). The analytic signal represents a narrowband signal in the time domain as a rotating vector, calculated as { *a*(*t*) = *A*(*t*). *cos*(*φ*(*t*)) }, where *A*(*t*) is the time-varying amplitude with carrier frequency expressed by the time-varying phase *φ*(*t*). The amplitude is determined by the modulus and the phase is determined by the argument of the complex signal, *z*(*t*), {*z*(*t*) = *a*(*t*) + *i.HT* [*a*(*t*)]}, where *HT* [*a*(*t*)] is the Hilbert transform of the analytical signal, *a*(*t*), and *i* is the imaginary unit (Glerean et al., 2012; Ponce-Alvarez et al., 2015; Deco et al., 2019a). Figure 1.1B shows the representation of the Hilbert BOLD phase for a brain area over time in the complex plane.

**Figure 1:**
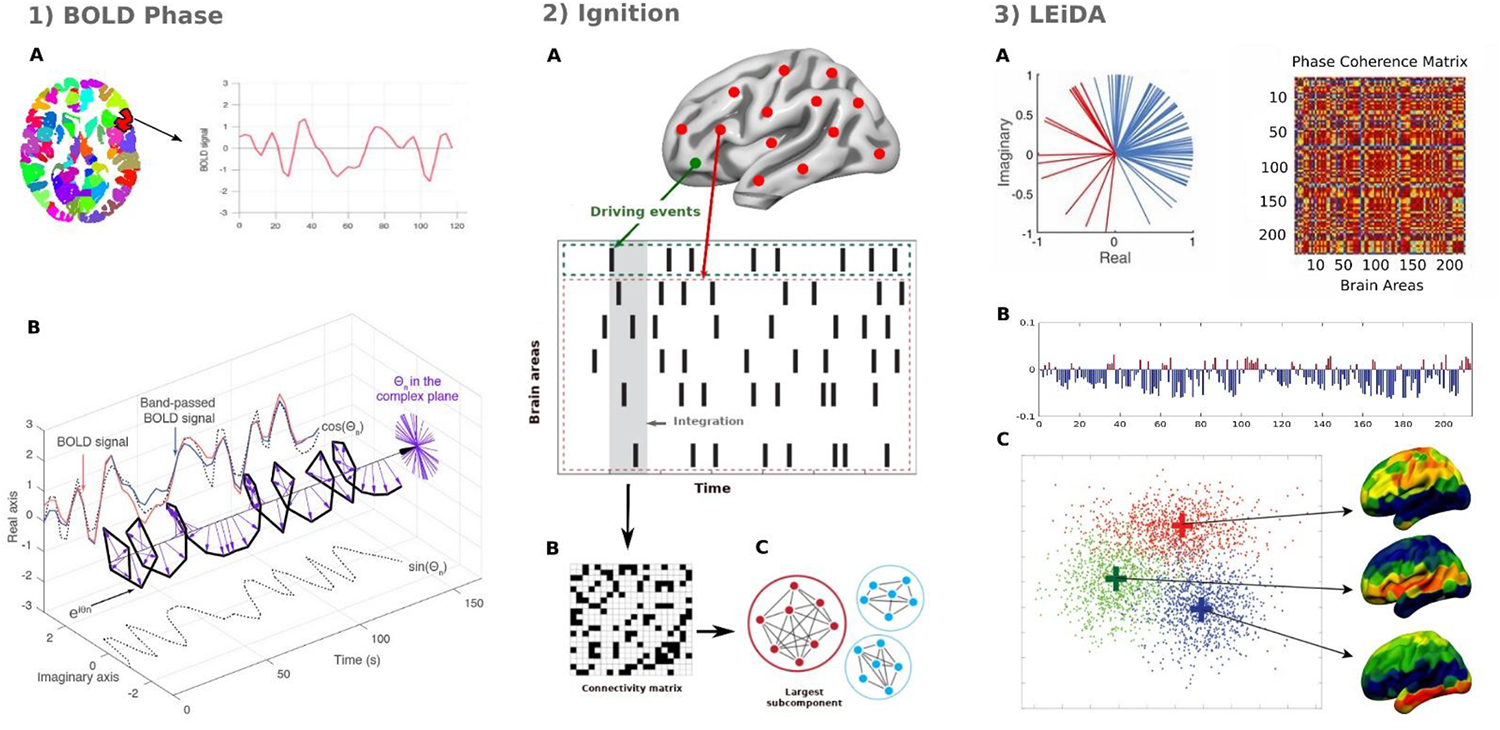
Methods. We applied two data-driven whole-brain methods based on phase synchronization of the BOLD signals. (1) For each of the 214 brain areas, we extracted the BOLD time series and computed the phase space of the BOLD signal. (1A) Specifically, we obtained the time series for each brain area using a resting-state atlas (Shen et al., 2013). (1B) Then, we measured the phase space of the BOLD signal by using the Hilbert transform for each brain area. The BOLD signal (red) was band-pass filtered between 0.04 and 0.07 Hz (blue) and converted with the Hilbert transform into an analytical signal represented by its instantaneous amplitude A(t) and its phase φ (with real and imaginary components). The phase dynamics can be represented in the complex plane as e^iφ^ (black bold line), the real part as cosφ (black dotted line), and the imaginary part as sinφ (black dotted line). The purple arrows represent the Hilbert phases at each TR. (2) Measuring intrinsic ignition. (2A) Events were captured by applying a threshold method (Tagliazucchi et al., 2012) (see green area). For each event evoked, the activity in the rest of the network (see red stippled area) was measured in the 4TR time window (gray area). (2B) A binarized phase lock matrix was obtained from the time window. (2C) From this phase lock matrix, we obtained the integration by calculating the largest subcomponent (i.e., by applying the global integration measure (Deco et al., 2015, 2017b)). Repeating the process for each driving event, we obtained the ignition and metastability of the intrinsic-driven integration for each brain area across the whole-brain network. (3) Finally, we applied the Leading Eigenvector Dynamics Analysis (LEiDA) to characterize differences between groups in dynamic functional connectivity patterns or metastable substates. (3A) The left panel shows the BOLD phases in all 214 brain areas represented in the complex plane. The right panel shows the phase coherence matrix between each pair of brain areas. (3B) The leading eigenvector V1(t) from this matrix was extracted. (3C) We applied a k-means clustering algorithm to detect the metastable substates from all the leading eigenvectors, across timepoints, number of subjects, and groups. Figure adapted from (Deco and Kringelbach, 2017, Deco et al., 2019a).

### Intrinsic-Ignition Framework

To measure the effect of spontaneous local activation events on whole-brain integration, we applied the Intrinsic-Ignition Framework (Deco and Kringelbach, 2017) using the phase space of the signals. This framework has been successfully applied in different resting-state fMRI studies (Deco et al., 2017b; Escrichs et al., 2019; Padilla et al., 2019; Alonso-Martínez et al., 2020). This approach characterizes the spatiotemporal propagation of information by measuring the degree of integration among spontaneous occurring events across the brain over time. Figure 1.2 represents the algorithm used to obtain the ignition value of each brain area evoked by an event within a set time window. Specifically, we averaged across the events the integration evoked at each time *t* with the time window set at 4TR. A binary event is defined by transforming the time series into z-scores, *z_i_*(*t*), and fixing a threshold, θ, given by the sum of the mean and the standard deviation of the signal in each brain area, such that the binary sequence σ(*t*) = 1 *z_i_*(*t*) > θ and crosses the threshold from below, and σ(*t*) = 0 otherwise (Figure 1.2A) (Tagliazucchi et al., 2012; Deco et al., 2017b). First, we obtained the instantaneous phase in all brain areas as explained in the Phase Synchronization section above and Figure 1.1. Then, we calculated the phase lock matrix *P_jk_*(*t*), which describes the state of pair-wise phase synchronization at time *t* between regions *j* and *k* as:

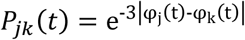

where φ_j_(t) and φ_k_(t) correspond to the obtained phase of the brain areas *j* and *k* at time *t*. Then, the integration is defined by measuring the length of largest connected component in the binarized symmetric phase lock matrix *P_jk_*(*t*) (Figure 1.2B). That is, given the fixed threshold θ, the matrix is binarized such that (0 if |*P_jk_*| < θ, 1 otherwise), and the integration value is computed as the length of the connected component considered as an adjacent graph (i.e., the largest subcomponent) (Figure 1.2C). The largest subcomponent represents the broadness of communication across the network for each driving event (Deco et al., 2015). Finally, repeating the process for each event in each brain area, the framework returns the mean integration and the standard deviation across the network. The mean integration is called ignition and it represents the spatial diversity; the standard deviation is called metastability, and it represents the variability over time for each brain area. Greater metastability in a brain area means that its activity changes more frequently across time within the network. The framework was computed across the whole-brain functional network (214 brain areas), as well as independently for eight resting-state networks: the frontoparietal, medial frontal, default-mode, subcortical, motor, visual I, visual II, and visual-association networks (Finn et al., 2015).

### Leading Eigenvector Dynamics Analysis (LEiDA)

To identify differences between groups in recurrent patterns of time-varying connectivity (dynamic functional connectivity) or ‘metastable-substates’ across all subjects, we used Leading Eigenvector Dynamics Analysis (LEiDA) (Cabral et al., 2017), a k-means clustering analysis based on the phase synchronization of BOLD signals. First, we computed a dynamic phase coherence connectivity matrix (Deco and Kringelbach, 2016) with size NxNxT, where N=214 (total number of brain areas), and T=117 (total number of timepoints), using the Hilbert transform as explained above in the Phase Synchronization section. Then, we calculated the BOLD phase coherence matrix (Figure 1.3A) at time *t* between each pair of brain areas *n* and *p* by computing the cosine of the phase difference as:

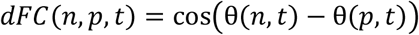

Given that the Hilbert transform expresses any signal in the polar coordinate system (i.e., *a*_(*t*)_ = *A*(*t*) · cos(φ_(*t*)_)), when the cosine function is applied, two brain areas *n* and *p* with similar angles at a given time *t* will show a phase coherence near 1 (i.e., *cos*(0°) = 1), whereas two brain areas that are orthogonal at a given time *t* will show a phase coherence near zero (i.e., *cos*(90°) = 0) (Cabral et al., 2017; Deco et al., 2019b). Second, to characterize the dynamic functional connectivity patterns across all subjects and timepoints, we obtained a leading eigenvector *V*_1_(*t*) for each *dFC*(*t*) at time *t* by capturing the dominant functional connectivity pattern rather than the whole set of matrices. This approach allows reducing the dimensionality of the data considerably because it only considers a single *V*_1_(*t*) for each dynamic functional connectivity matrix. The *V*_1_(*t*) is an Nx1 vector capturing the principal orientation of the BOLD phase (showing positive or negative values) for each of the 214 brain areas (Figure 1.3B). Finally, we applied a k-means clustering algorithm using a range from k=2 to k=7 clusters to detect metastable substates or dynamic functional connectivity states from all the leading eigenvectors *V*_1_(*t*) across timepoints, subjects, and groups: 117 timepoints x 310 subjects x 2 groups = 72,540 *V*_1_(*t*). We obtained k cluster centroids, each one as an Nx1 vector, which represent recurrent metastable substates across all subjects. The clustering configuration that best represented the resting-state data of all 620 subjects and distinguished between the two groups was detected at *k* = 3 (Figure 1.3C). We rendered the resulting cluster centroids onto a surface cortex using Surf Ice (https://www.nitrc.org/projects/surfice/). A complete description of the method can be found in Cabral et al. (2017).

### Statistical analysis

Statistical analyses were done with software MATLAB version R2017a (MathWorks, Natick, MA, USA). We applied a Monte Carlo permutation method to test the results of the Intrinsic-Ignition Framework (ignition and metastability) and to test the results of the LEiDA method (probability of occurrence and duration of each metastable substate). More specifically, we randomly shuffled the labels for each pair of conditions to be tested and created two new simulated conditions (10,000 iterations). Then, we measured how many times the difference between the new simulated conditions was greater than the difference between the real conditions; in other words, we calculated the p-value of the null hypothesis that the two random distributions show a greater difference than the real conditions. Furthermore, we applied the False Discovery Rate (FDR) method (Hochberg and Benjamini, 1990) to correct for multiple comparisons when necessary.

## Results

### Intrinsic Ignition

We computed the Intrinsic-Ignition Framework across the whole-brain functional network and found that the mean ignition was higher in the older group than in the middle-age group (p<0.001) (Figure 2a). In the middle-age group, the regions with the highest intrinsic ignition belong to the visual networks, subcortical network, frontoparietal network, motor network, and medial-frontal network: the right middle occipital gyrus, right middle temporal gyrus, right lingual gyrus, fusiform gyri, left hippocampus and parahippocampal gyrus, right inferior temporal gyrus, right superior temporal gyrus, left calcarine fissure and surrounding cortex, left precentral gyrus, and right insula. By contrast, in the older group, the regions showing the highest intrinsic ignition areas belong to the visual networks, medial frontal network, and frontoparietal network: the right middle occipital, middle temporal gyrus, left fusiform gyrus, lingual gyrus, right inferior temporal gyrus, right middle frontal gyrus, calcarine fissure and surrounding cortex in both hemispheres, left superior frontal gyrus, left inferior frontal gyrus, left insula, left thalamus, and right cuneus. Table 2 shows the 20 brain areas with the highest intrinsic-ignition capability for each group.

**Figure 2:**
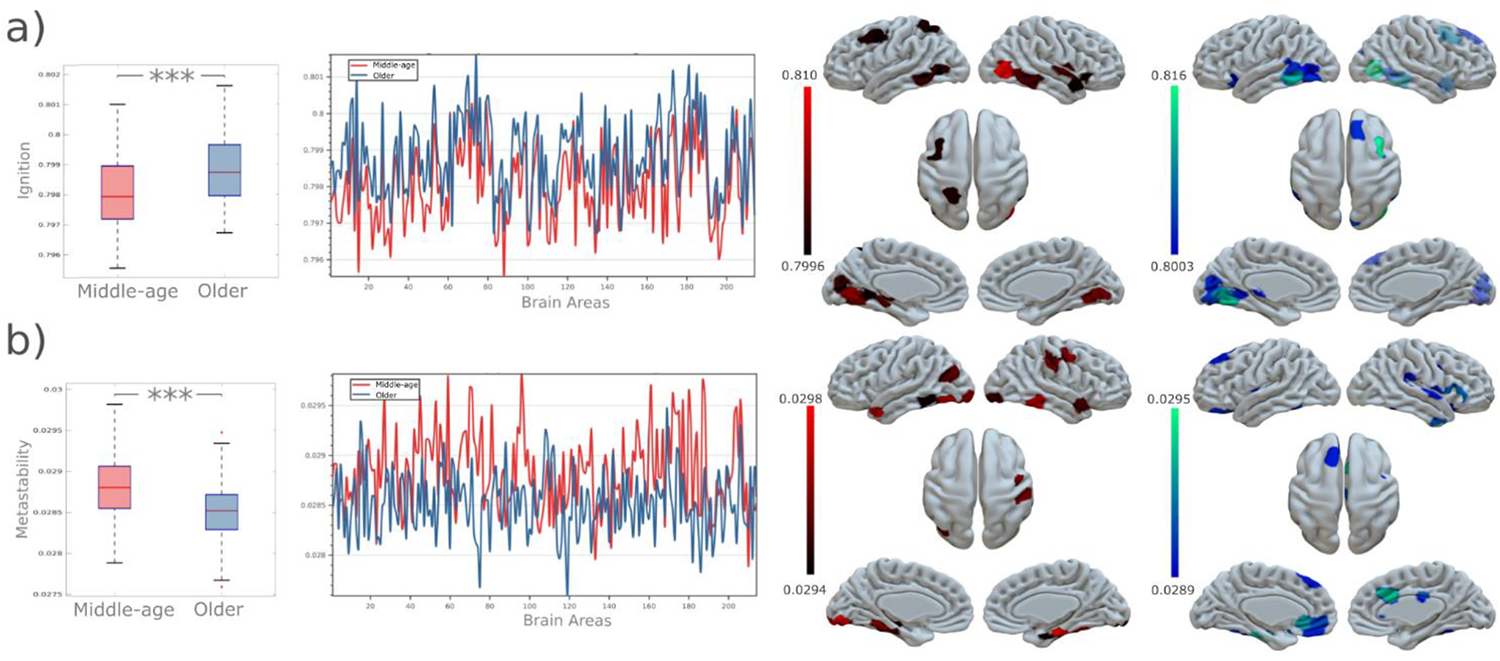
Intrinsic Ignition Framework. (a) Ignition measure. The boxplot shows the mean integration (ignition) for each group (middle-age group and older group). The ignition was higher in the older group (in blue) than in the middle-age group (in red) (p < 0.001). The second graph shows ignition plotted across brain areas. Rendered brains represent the 20 regions with the highest ignition for each group (middle-age in red and older group in blue). (b) Metastability measure. By contrast, the middle-age group showed higher metastability across the whole-brain compared to the older group (p < 0.001). P-values are based on Monte-Carlo permutation tests, *** represents p<0.001.

**Table 2.** Intrinsic ignition capability

| Middle-age group |  |  | Older Group |  |  |
| --- | --- | --- | --- | --- | --- |
| rs-fMRI atlas | Corresponding AAL-regions | Network | rs-fMRI atlas | Corresponding AAL-regions | Network |
| 74 | 34% Middle occipital gyrus, right [52]<br>32% Middle temporal gyrus, right [86] | Visual association | 74 | 34% Middle occipital gyrus, right [52]<br>32% Middle temporal gyrus, right [86] | Visual association |
| 185 | 88% Lingual gyrus, left [47] | Visual | 181 | 40% Fusiform gyrus, left [55]<br>31% Lingual gyrus, left [47] | Visual_I |
| 79 | 78% Lingual gyrus, right [48] | Visual | 173 | 63% Inferior temporal gyrus, left [89] | Frontoparietal |
| 72 | 51% Lingual gyrus, right [48]<br>33% Fusiform gyrus, right [56] | Visual | 185 | 88% Lingual gyrus, left [47] | Visual_I |
| 181 | 40% Fusiform gyrus, left [55]<br>31% Lingual gyrus, left [47] | Visual | 70 | 70% Inferior temporal gyrus, right [90] | Frontoparietal |
| 205 | 43% Parahippocampal gyrus, left [39]<br>24% Hippocampus, left [37] | Subcortical | 14 | 85% Middle frontal gyrus, right [8] | Frontoparietal |
| 69 | 51% Inferior temporal gyrus, right [90]<br>47% Middle temporal gyrus, right [86] | Visual association | 53 | 45% Temporal pole: superior, right [84]<br>34% Temporal pole: middle temporal, right [88] | Medialfrontal |
| 70 | 70% Inferior temporal gyrus, right [90] | Frontoparietal | 69 | 51% Inferior temporal gyrus, right [90]<br>47% Middle temporal gyrus, right [86] | Visual association |
| 61 | 50% Superior temporal gyrus, right [82]<br>28% Rolandic operculum, right [18] | Motor | 187 | 38% Lingual gyrus, left [47]<br>20% Calcarine fissure and surrounding cortex, left [43] | Visual_II |
| 189 | 67% Calcarine fissure and surrounding cortex, left [43] | Visual_I | 127 | 46% Inferior frontal gyrus, orbital part, left [15]<br>36% Insula, left [29] | Medialfrontal |
| 173 | 63% Inferior temporal gyrus, left [89] | Frontoparietal | 166 | 82% Middle temporal gyrus, left [85] | Medialfrontal |
| 183 | 41% Middle temporal gyrus, left [85]<br>35% Middle occipital gyrus, left [51] | Visual association | 180 | 49% Inferior occipital gyrus, left [53]<br>33% Fusiform gyrus, left [55] | Visual association |
| 36 | 52% Insula, right [30]<br>36% Inferior frontal gyrus, orbital, right [16] | Subcortical | 189 | 67% Calcarine fissure and surrounding cortex, left [43] | Visual_I |
| 149 | 66% Superior parietal gyrus, left [59] | Visual association | 80 | 39% Calcarine fissure and surrounding cortex, right [44]<br>27% Cuneus, right [46] | Visual_I |
| 123 | 84% Middle frontal gyrus, left [7] | Medial frontal | 76 | 48% Lingual gyrus, right [48]<br>17% Fusiform gyrus, right [56] | Visual_II |
| 139 | 81% Precentral gyrus, left [1] | Medial frontal | 12 | 52% Superior frontal gyrus, dorsolateral, right [4]<br>41% Superior frontal gyrus, medial, right [24] | Medialfrontal |
| 179 | 57% Lingual gyrus, left [47]<br>21% Calcarine fissure and surrounding cortex, left [43] | Visual_I | 179 | 57% Lingual gyrus, left [47]<br>21% Calcarine fissure and surrounding cortex, left [43] | Visual_I |
| 53 | 45% Temporal pole: superior, right [84]<br>34% Temporal pole: middle temporal, right [88] | Medial frontal | 212 | 46% Thalamus, left [77]<br>1% Lingual gyrus, left [47] | Subcortical |
| 162 | 54% Temporal pole: superior temporal gyrus, left [83]<br>27% Superior temporal gyrus, left [81] | Motor | 183 | 41% Middle temporal gyrus, left [85]<br>35% Middle occipital gyrus, left [51] | Visual association |
| 63 | 52% Superior temporal gyrus, right [82]<br>48% Middle temporal gyrus, right [86] | Motor | 79 | 78% Lingual gyrus, right [48] | Visual_I |

AAL: Anatomical Automatic Labeling Atlas

Metastability was lower in the older group than in the middle-age group (p<0.001) (Figure 2b). In the middle-age group, the brain areas with the highest metastability belong mainly to the default-mode network, visual networks, motor network, and frontoparietal network: the parahippocampal gyri, fusiform gyri, left inferior temporal gyrus, left lingual gyrus, left hippocampus, middle temporal gyri, right inferior occipital gyrus, right precentral gyrus, and right postcentral gyrus. By contrast, in the older group, the brain areas with the highest metastability belong mainly to the motor network, subcortical network, default-mode network, medial frontal network, and visual association network: the inferior temporal gyri, left fusiform gyrus, left superior frontal gyrus, inferior frontal gyrus, right anterior cingulate and paracingulate gyri, right median cingulate gyrus, bilateral insula, right superior temporal gyrus, left rectus gyrus, bilateral Rolandic opercula, left parahippocampal gyrus, and right precentral gyrus. Table 3 shows the 20 brain areas with the highest metastability for each group.

**Table 3.** Metastability

| Middle-age group |  |  | Older Group |  |  |
| --- | --- | --- | --- | --- | --- |
| rs-fMRI atlas | Corresponding AAL-regions | Network | rs-fMRI atlas | Corresponding AAL-regions | Network |
| 96 | 60% Parahippocampal gyrus, right [40]<br>28% Fusiform gyrus, right [56] | Default mode | 169 | 46% Inferior temporal gyrus, left [89]<br>41% Fusiform gyrus, left [55] | Motor |
| 59 | 55% Fusiform gyrus, right [56]<br>41% Inferior temporal gyrus, right [90] | Visual association | 15 | 56% Median cingulate and paracingulate gyri, right [34]<br>26% Anterior cingulate and paracingulate gyri, right [32] | Subcortical |
| 187 | 38% Lingual gyrus, left [47]<br>20% Calcarine fissure and surrounding cortex [43] | Visual_II | 206 | 55% Fusiform gyrus, left [55]<br>32% Parahippocampal gyrus, left [39] | Subcortical |
| 161 | 57% Temporal pole: middle temporal gyrus, left [87]<br>25% Middle temporal gyrus, left [85] | Medial frontal | 108 | 33% Anterior cingulate and paracingulate gyri, left [31]<br>23% Rectus gyrus, left [27] | Default mode |
| 70 | 70% Inferior temporal gyrus, right [90] | Frontoparietal | 109 | 51% Inferior frontal gyrus, orbital part, left [15]<br>31% Superior frontal gyrus, orbital part, left [5] | Subcortical |
| 71 | 48% Fusiform gyrus, right [56]<br>29% Inferior temporal gyrus, right [90] | Visual association | 16 | 45% Inferior frontal gyrus, triangular part, right [14]<br>28% Inferior frontal gyrus, orbital, right [16] | Medialfrontal |
| 180 | 49% Inferior occipital gyrus, left [53]<br>33% Fusiform gyrus, left [55] | Visual association | 88 | 42% Median cingulate and paracingulate gyri, right [34] | Subcortical |
| 188 | 25% Inferior occipital gyrus, left [53]<br>23% Lingual gyrus, left [47] | Visual_II | 57 | 67% Inferior temporal gyrus, right [90] | Medialfrontal |
| 172 | 71% Fusiform gyrus, left [55] | Visual_I | 18 | 59% Inferior frontal gyrus, orbital, right [16]<br>20% Insula, right [30] | Subcortical |
| 27 | 81% Precentral gyrus, right [2] | Motor | 112 | 54% Superior frontal gyrus, medial orbital, left [25]<br>31% Anterior cingulate and paracingulate gyri, left [31] | Default mode |
| 45 | 47% Supramarginal gyrus, right [64]<br>35% Postcentral gyrus, right [58] | Motor | 174 | 58% Fusiform gyrus, left [55]<br>36% Inferior temporal gyrus, left [89] | Visual association |
| 206 | 55% Fusiform gyrus, left [55]<br>32% Parahippocampal gyrus, left [39] | Subcortical | 142 | 59% Insula, left [29]<br>23% Rolandic operculum, left [17] | Motor |
| 177 | 73% Middle occipital gyrus, left [51] | Default mode | 175 | 65% Inferior temporal gyrus, left [89] | Visual association |
| 53 | 45% Temporal pole: superior, right [84]<br>34% Temporal pole: middle temporal, right [88] | Medial frontal | 111 | 62% Rectus gyrus, left [27] | Medial frontal |
| 81 | 59% Inferior occipital gyrus, right [54]<br>23% Lingual gyrus, right [48] | Visual_II | 122 | 44% Superior frontal gyrus, medial, left [23]<br>43% Superior frontal gyrus, dorsolateral, left [3] | Medial frontal |
| 204 | 46% Hippocampus, left [37]<br>19% Inferior temporal gyrus, left [89] | Subcortical | 46 | 53% Superior temporal gyrus, right [82]<br>37% Supramarginal gyrus, right [64] | Motor |
| 175 | 65% Inferior temporal gyrus, left [89] | Visual association | 71 | 48% Fusiform gyrus, right [56]<br>29% Inferior temporal gyrus, right [90] | Visual association |
| 173 | 63% Inferior temporal gyrus, left [89] | Frontoparietal | 61 | 50% Superior temporal gyrus, right [82]<br>28% Rolandic operculum, right [18] | Motor |
| 97 | 67% Parahippocampal gyrus, right [40] | Motor | 31 | 52% Precentral gyrus, right [2]<br>22% Inferior frontal gyrus, opercular part, right [12] | Frontoparietal |
| 203 | 55% Hippocampus, left [37]<br>13% Parahippocampal gyrus, left [39] | Subcortical | 55 | 67% Inferior temporal gyrus, right [90] | Frontoparietal |

**Table 4.**
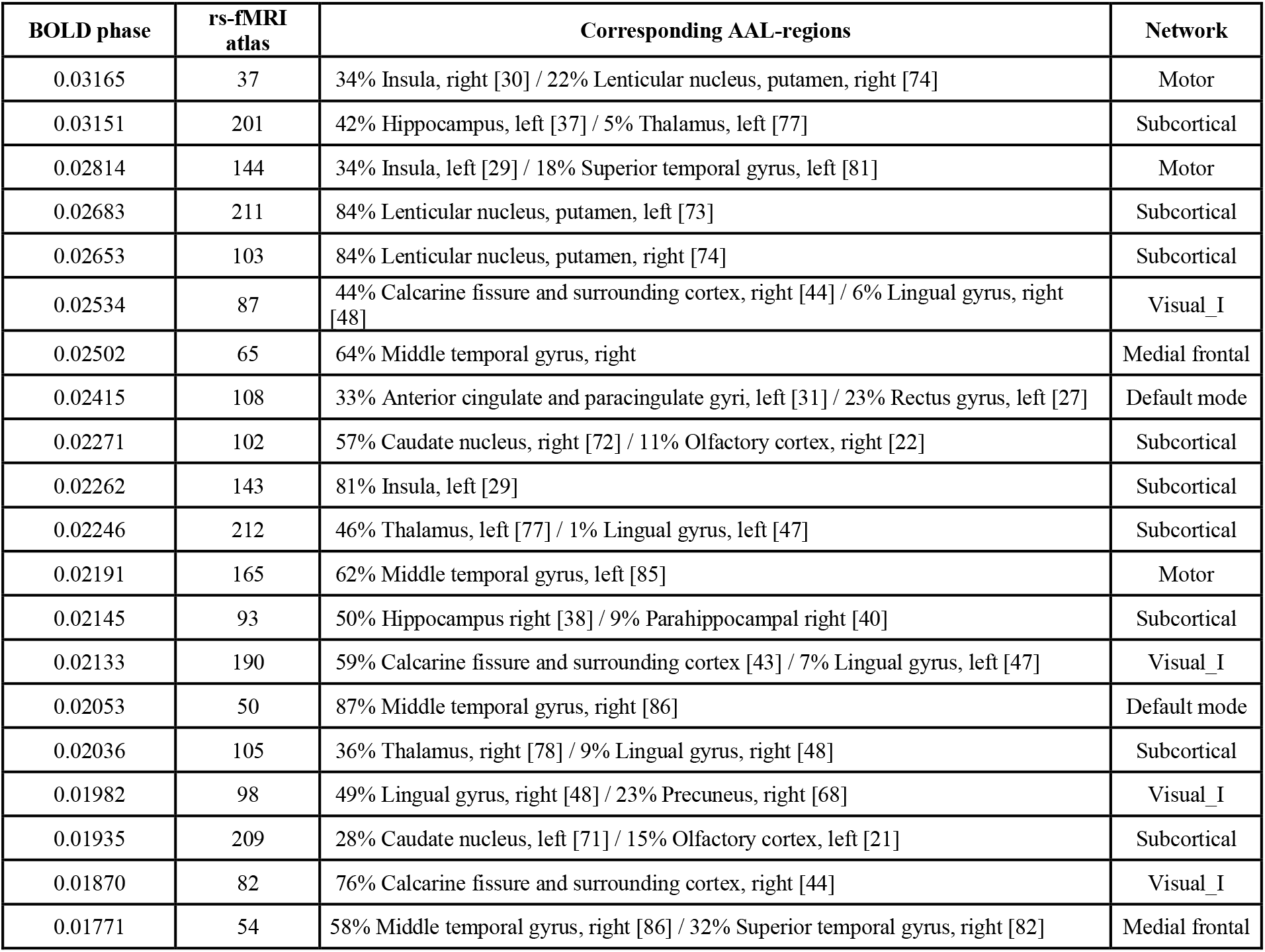
Cluster centroids of the significant metastable substate

Moreover, we computed the intrinsic ignition and metastability independently for each resting-state network. Figure 3 shows the absolute difference between the middle-age and older groups in the intrinsic-ignition values for each brain area in each network. Compared to the middle-age group, the older group had significantly increased intrinsic ignition in the frontoparietal network (FDR-corrected, p<0.001) and medial frontal network (FDR-corrected, p<0.001). By contrast, the middle-age group had greater intrinsic ignition in the motor network (FDR-corrected, p<0.001). There were no significant differences between groups in intrinsic ignition in the default-mode, subcortical, visual I, visual II, or visual-association networks. Figure 4 shows the absolute difference between the middle-age and older groups in metastability values for each brain area in each network. Compared to the middle-age group, the older group had significantly increased metastability in the frontoparietal network (FDR-corrected, p<0.01) and medial frontal network (FDR-corrected, p<0.01). By contrast, the middle-age group had greater metastability in the defaultmode (FDR-corrected, p<0.05), subcortical (FDR-corrected, p<0.001), motor (FDR-corrected, p<0.001), visual association (FDR-corrected, p<0.05), and visual I networks (FDR-corrected, p<0.001). Only the visual II network did not differ significantly between groups.

**Figure 3:**
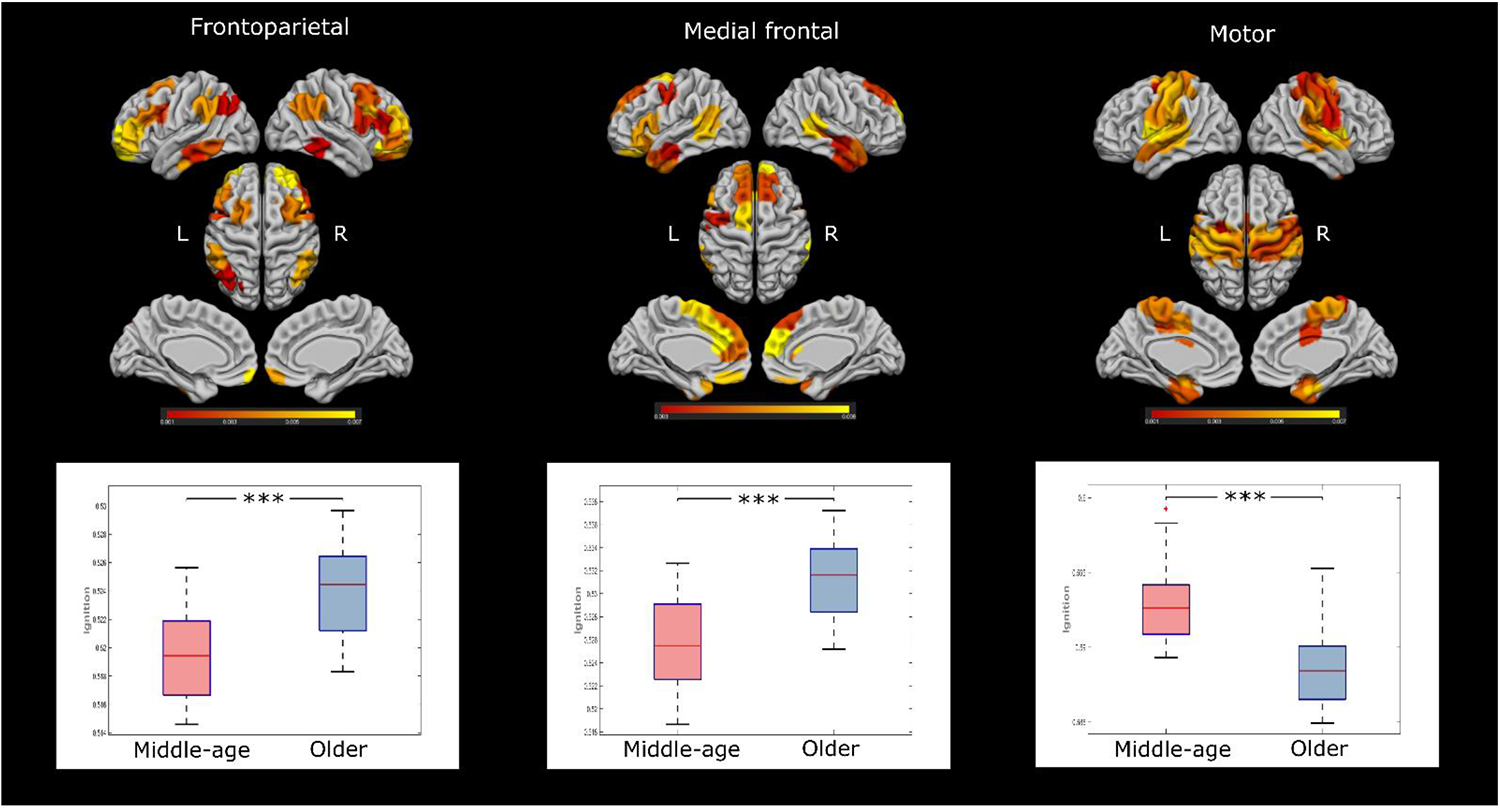
Differences in ignition across resting-state networks. The plots show the differences between groups in each significant resting-state network; rendered brains represent the absolute difference in ignition values for each brain area in each network between the middle-age and older groups (the greatest difference is marked in yellow). Compared to the middle-age group, intrinsic ignition was significantly higher in the older group in the frontoparietal network (FDR-corrected, p<0.001) and medial frontal network (FDR-corrected, p<0.001). By contrast, intrinsic ignition was significantly higher in the middle-age group in the motor network (FDR-corrected, p<0.001). The default-mode, subcortical, visual I, visual II, and visual association networks were not significantly different between groups.

**Figure 4:**
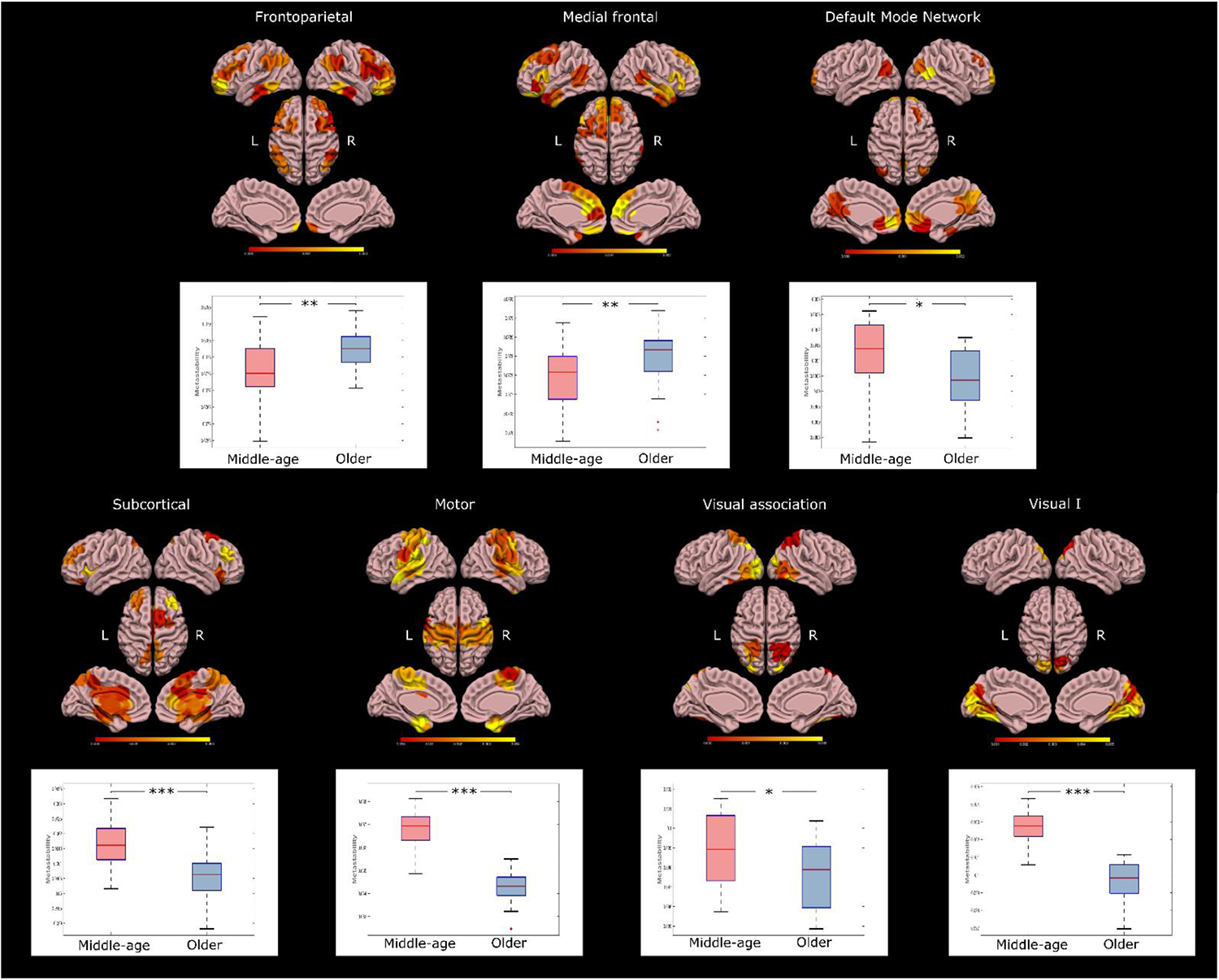
Differences in metastability across resting-state networks. The plots show the differences between groups in each significant resting-state network, whereas rendered brains represent the absolute difference (middle-age and older) between metastability values for each brain area in each network (in yellow the highest difference). The older group showed significantly increased metastability compared to the middle-age group in the frontoparietal network (FDR-corrected, p<0.01) and medial frontal network (FDR-corrected, p<0.01), whereas the middle-age group showed increased metastability in the default-mode network (FDR-corrected, p<0.05), subcortical network (FDR-corrected, p<0.001), motor network (FDR-corrected, p<0.001), visual association network (FDR-corrected, p<0.05), and visual I network (FDR-corrected, p<0.001).

### LEiDA

Clustering across all subjects and timepoints identified three metastable substates. Figure 5A compares the probability of occurrence of each metastable substate between groups, and Figure 5B compares the duration of these substates between groups. Figure 5C shows the three metastable substates rendered onto a surface cortex. The metastable substate that had the highest probability of occurrence (the first metastable substate) closely overlaps with the state of global BOLD coherence (Cabral et al., 2017). The probability of this substate occurring was higher in the older group [0.476 ± 0.008 (mean ± standard error) vs. 0.453 ± 0.008 in the middle-age group, FDR-corrected *p* = 0.03], and this substate also lasted longer in the older age group [32.465 ± 0.957 seconds vs. 30.265 ± 0.791 seconds in the middle-age group, *p* = 0.04], although the difference in duration was no longer significant after FDR correction. The second metastable substate is especially interesting because it closely overlaps with the so-called rich club (Hagmann et al., 2008; van den Heuvel and Sporns, 2011; van den Heuvel et al., 2012; Sporns, 2013). In particular, this substate involved the following areas in both hemispheres: the superior frontal cortex, precuneus, insula, and subcortical areas, such as the caudate, putamen, hippocampus, and thalamus (see Figure 5D). The networks most frequently involved in this metastable substate were the subcortical network, visual network, motor network, default-mode network, and medial frontal network. The probability of this substate occurring was greater in the middle-age group [0.288 ± 0.007 vs. 0.269 ± 0.006 in the older group, FDR-corrected *p* = 0.026], and this substate also lasted longer in the middle-age group [16.399 ± 0.605 seconds vs. 14.853 ± 0.414 seconds in the older group, FDR-corrected *p* = 0.01). The third metastable substate was not significantly different between groups in its probability of occurrence (*p* = 0.35) or duration (*p* = 0.39).

**Figure 5:**
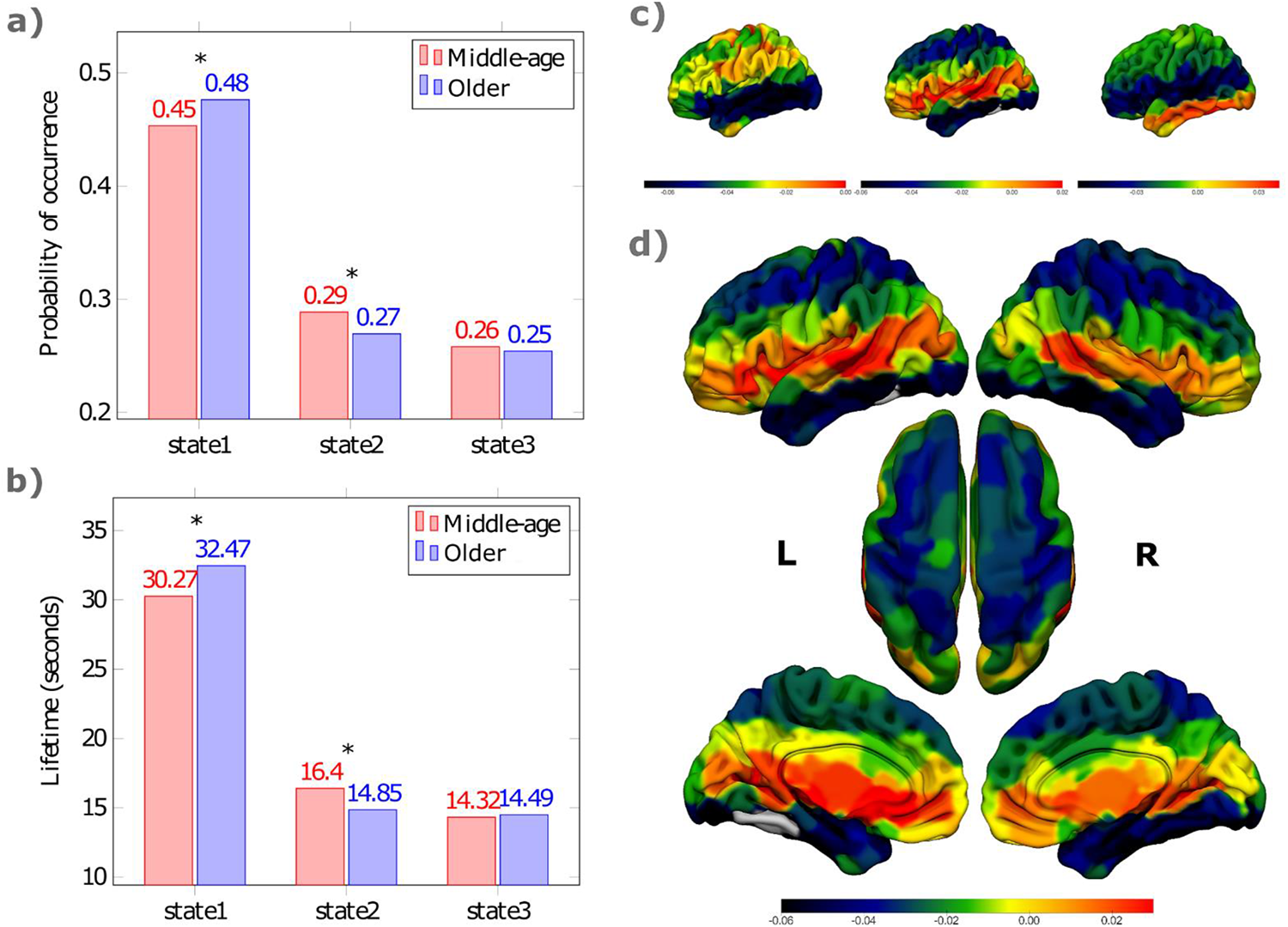
Metastable substates obtained by clustering. We identified three metastable substates that occurred frequently across all subjects during rest. The states are shown from higher to lower probability of occurrence. A) Differences in probability and B) differences in duration of each metastable substate between the middle-age group (in red) and the older group (in blue) during the resting-state scan. C) Metastable substates rendered on the cortex with Surfice. The first metastable substate had the highest probability of occurrence. This state was more likely to occur in subjects in the older group (mean, 0:476 ± 0:008 (s.e.) vs.0:453 ± 0:008 in the middle-age group, FDR-corrected p = 0.03); the duration of this state was also higher in the older group, although this comparison was no significant after FDR correction (32:465 ± 0:957 s vs. 30:265 ± 0:791 in the middle-age group, p = 0.04, uncorrected). The second metastable substate is especially interesting because it overlaps with the rich club. The probability of this state occurring was higher in the middle-age group (mean 0:288 ± 0:007 (s.e.) vs. 0:269 ± 0:006 in the older group, FDR-corrected p = 0.026); the duration of this state was also higher in the middle-age group (mean 16:399 ± 0:605 s vs. 14:853 ± 0:414 s in the older group, FDR-corrected p = 0.01). The third metastable substate was not significantly difference between groups in its probability of occurrence (p = 0.35) or duration (p = 0.39). D) Relevant metastable substate overlapping with rich-club regions in both hemispheres (the superior frontal cortex, precuneus, insula and subcortical areas such as the caudate, putamen, hippocampus, and thalamus).

## Discussion

Interest in characterizing resting-state functional patterns during aging is growing. Understanding the underlying dynamics across the whole-brain functional network may help us better understand age-related changes. In this line, various methods have been developed to capture statistical properties of resting-state fMRI data beyond classical static functional connectivity, providing a new perspective to interpret brain functioning during the resting scan. To investigate the underlying whole-brain dynamics, we applied two data-driven whole-brain methods based on phase coherence synchronization (Deco and Kringelbach, 2017; Cabral et al., 2017) to compare intrinsic ignition, metastability, and metastable substates between middle-aged and older subjects from a large sample of healthy human adults. To characterize the spatiotemporal propagation of information, we used the Intrinsic-Ignition Framework to measure the degree of integration of spontaneously occurring events across the whole-brain during rest. Ignition values across the whole-brain functional network were higher in older subjects than in middle-aged subjects, but older subjects also had less metastability. Applying Leading Eigenvector Dynamics Analysis (LEiDA), we found differences between groups in the probability of occurrence and duration of a metastable substate involving rich-club brain areas.

Interestingly, the older group had higher intrinsic ignition across the whole-brain functional network (Figure 2a); the brain areas with the highest intrinsic-ignition values were mainly distributed across the visual networks, frontoparietal network, and medial frontal network (Figure 2a and Table 2). The mean intrinsic-ignition value reflects spatial diversity and the broadness of communication across the whole network. These results are in line with previous studies investigating the effects of aging in resting-state networks. Geerligs et al. (2015) reported increased connectivity in older adults between the visual network and somatomotor network as well as between the visual network and cingulo-opercular network. Betzel et al. (2014) found increased functional connectivity between the dorsal attention network and the salience/ventral attention networks in older adults. Similarly, Spreng et al. (2016) found increased between-network functional connectivity across the default-mode network and dorsal attention networks during both task and rest conditions. We conclude that increased functional connectivity between resting-state networks has a significant impact across the whole-brain functional network as evidenced by the level of intrinsic ignition, and that the higher intrinsic ignition in the older group may be related to compensatory mechanisms.

Metastability was higher in the middle-age group (Figure 2b and Table 3). This finding is particularly interesting because middle-age adults showed lower intrinsic ignition across the whole-brain functional network compared to older adults, but the underlying dynamics of the middle-age adults seem to be more complex across time. Metastability characterizes the hierarchy of information processing in the brain. Thus, brain areas showing higher metastability are more relevant for the broadcasting of information than those showing lower. Greater metastability also reflects more complex brain dynamics (i.e., a more flexible switching across time), whereas lower metastability suggests a more stable system (Deco and Kringelbach, 2017; Deco et al., 2017a; Jobst et al., 2017). Our findings are in line with previous studies on the effects of aging on brain functional dynamics. For example, the decreased metastability in the older group in our study echoes recent studies that suggest deficient network modulation in the elderly (Turner and Spreng, 2015; Damoiseaux, 2017). Xia et al. (2019) found that the number of transitions between different metastable substates decreased with age, leading them to conclude that resting mind states may shift faster in young people than in older people. Similarly, variability across large-scale networks decreases linearly with aging over the lifespan (Nomi et al., 2017) and in healthy elderly subjects (Lou et al., 2019). Moreover, our findings that areas in the temporal and occipital regions were the most important for the broadcasting of information in the middle-age group (Figure 2b and Table 3) is consistent with the results of recent time-varying resting-state fMRI studies (Nomi et al., 2017; Kumral et al., 2019). Similarly, our findings that the frontal and temporal areas were more relevant in the older group (Figure 2b and Table 3) are consistent with the results of a recent EEG study that found an enhanced brain dynamics of phase synchronization in the alpha-band frequency, predominantly in frontal areas (Nobukawa et al., 2019), which the authors suggest could reflect a general change in functional connectivity dynamics during aging. Moreover, overactivation in prefrontal brain areas has been previously observed in older adults during fMRI tasks, giving rise to different theories (Cabeza, 2002; Davis et al., 2008; Reuter-Lorenz and Cappell, 2008).

We also explored intrinsic ignition and metastability across large-scale networks, computing the intrinsic-ignition framework within eight resting-state networks. In the older group, the frontoparietal and medial frontal networks showed higher ignition and metastability (Figures 3 and 4). These findings are in line with those reported by Lou et al. (2019), who found that the frontal and temporal lobes show a more dynamic pattern with increasing age. A recent meta-analysis pointed out that age-related changes in activation commonly affect the frontoparietal and default-mode networks (Li et al., 2015). The frontoparietal network serves as a flexible hub and plays a vital role in adaptive control and implementation of different responses to demands during tasks (Cole et al., 2013). The frontoparietal network is also involved in selecting relevant information from the environment (Ptak, 2012). The default-mode and frontoparietal networks are also thought to be critical in controlling global brain dynamics (Hellyer et al., 2014).

In the present study, metastability within the default-mode, subcortical, and visual-association networks was higher in the middle-age group (Figure 4). In a recent study in a large cohort of young subjects, Lee et al. (2019) reported higher metastability in lower-order resting-state networks, such as the visual network and auditory network, which are involved in specialized, mostly externally driven functions. These networks’ greater metastability might reflect a greater capacity to change their functional configuration in response to diverse, rapidly changing external inputs (Power et al., 2011). By contrast, higher-order networks such as the default-mode and central executive networks are mostly involved in internal and goal-directed processing (Raichle et al., 2001; Raichle and Snyder, 2007), so it would make sense for their functional configurations to last longer. Moreover, the previously mentioned study also found that metastability was strongly associated with various indicators of higher-order cognitive ability and physical wellbeing (Lee et al., 2019).

One of the most noteworthy results in our study was the identification of a metastable substate overlapping the so-called the ‘rich club’ of densely interconnected nodes (Hagmann et al., 2008; van den Heuvel and Sporns, 2011; van den Heuvel et al., 2012; Sporns, 2013; Deco et al., 2017a). This substate involved the superior frontal cortex, precuneus, insula, and subcortical areas (caudate, putamen, hippocampus, and thalamus) in both hemispheres. It is thought that the rich club might also act as a gatekeeper that coordinates interactions with lower-degree regions and the emergence of different functional network configurations (van den Heuvel and Sporns, 2011). We found that the metastable substate corresponding to the rich club was less likely to occur in the older group and that when it did occur, it did so for shorter periods of time. Damoiseaux (2017) suggested that less-efficient rich-club network might be responsible for the differences in brain dynamics observed in older subjects. Our findings are in line with the hypothesis that the rich club connects different functional modules in the brain that partially overlap with different resting-state networks (Biswal et al., 1995; van den Heuvel and Sporns, 2011). Our findings regarding the lower probability of occurrence and shorter duration of this substate in the older group might be due to alterations in the intrinsic dynamics of this particular metastable substate or in any of the brain areas involved. Rich-club regions play a key role in integrating information across the brain network; consequently, damage to a brain area belonging to the rich club can affect global communication and have repercussions in multiple cognitive domains (van den Heuvel and Hulshoff Pol, 2010; Baggio et al., 2015; Deco and Kringelbach, 2017). Our results are consistent with the observation that the efficiency of the rich-club network increases during brain development in early life and decreases late in life in a manner that yields an inverted-U when plotted along the lifespan (Cao et al., 2014; Zhao et al., 2015; Damoiseaux, 2017).

Our LEiDA analysis also found that the first metastable substate, which has been related to the global signal in fMRI studies, had a higher probability of occurrence, and longer duration in the older group (although this last comparison was no longer significant after correction for multiple comparisons) (Figure 3). Like in previous resting-state fMRI studies applying LEiDA (Cabral et al., 2017; Figueroa et al., 2019; Lord et al., 2019), this anti-correlated state of global BOLD phase coherence (i.e., all BOLD phases showing negative values in the leading eigenvector) was the most prevalent. Although the significance of the global signal remains controversial, growing evidence suggests that it could contain valuable neurophysiological information and should not therefore be treated as a nuisance term (Saad et al., 2012; Liu et al., 2017). In a study with simultaneous fMRI and EEG acquisition during rest, Wong et al. (2013) found that increased EEG vigilance induced with caffeine was associated with decreased global signal amplitude and increased anti-correlation between the default-mode network and the task-positive network. Moreover, the global signal amplitude seems to increase during early sleep stages (Fukunaga et al., 2006). However, the role of the global BOLD phase coherence state remains unclear and needs further investigation (Cabral et al., 2017).

This study has several limitations. Although this cross-sectional study analyzed data from a large sample of healthy human adults, it would be very instructive to explore the age-related changes in neuroimaging in the same subjects in a longitudinal study. Data-driven methods alone are insufficient to understand the mechanisms underlying the process of aging or explain the causes of the dynamic changes observed. On the other hand, brain models simulating time series have advanced our understanding of the relationship between structure and function in the brain and the potential repercussions of disrupted connectivity from injury or disease; moreover, in silico simulations open the possibility of discovering potential stimulation targets to shift patients’ global brain dynamics toward a healthier state (Deco and Kringelbach, 2014; Deco et al., 2019a). One line for future studies could focus on assessing the behavioral relevance of intrinsic ignition and metastability through the aging process. Finally, although age is strongly associated with changes in functional connectivity, more studies are needed to further characterize brain functional connectivity in older adults and resolve inconsistent results due to methodological differences among studies.

In conclusion, applying two novel data-driven approaches to examine whole-brain dynamic changes, this work provides new insights into age-related brain changes. Our findings suggest that, compared to middle-aged subjects, older subjects show higher ignition but lower metastability across the whole-brain network, as well as reduced access to a dynamic functional connectivity pattern that is key for communication in the brain. These findings support the hypothesis that cognitive processing methods differ between middle-aged and older adults. Taken together, these findings suggest that functional whole-brain dynamics are altered in aging, probably due to an imbalance in a metastable substate that involves brain areas of the so-called rich club. Further investigations will surely improve our understanding of brain changes during aging.

## Acknowledgments

The Aging Imageomics Study was funded by the Government of Catalonia’s Department of Health’s *Pla Estratègic de Recerca i Innovació en Salut* (PERIS) 2016-2020 (file number, SLT002/16/00250). We also acknowledge funding from the Spanish Ministry of Science, Innovation, and Universities (RTI2018-099200-B-I00, co-financed by FEDER funds from the European Union (“A way to build Europe”)), and the Generalitat of Catalonia (2017SGR696) to RP. IRBLleida is a CERCA Programme/Generalitat of Catalonia. Toshiba Medical Systems (now Canon Medical Systems) provided a dedicated 1.5T MRI scanner and ancillary MRI equipment for this study. We would like to express our sincere gratitude to the subjects who participated in the Aging Imageomics Study for their valuable contribution and the study staff for coordination and data collection.

